# Midkine-a is required for cell cycle progression of Müller glia during neuronal regeneration

**DOI:** 10.1101/668210

**Authors:** Mikiko Nagashima, Travis S. D’Cruz, Doneen Hesse, Christopher J. Sifuentes, Pamela A. Raymond, Peter F. Hitchcock

## Abstract

In zebrafish, Müller glia function as intrinsic retinal stem cells that can regenerate ablated neurons. Understanding the mechanisms governing neuronal stem cells may provide clues to regenerate neurons in mammals. We report that in Müller glia the cytokine/growth factor, Midkine-a, functions as a core autocrine regulator of the cell cycle. Utilizing *midkine-a* mutants, we determined that Midkine-a regulates elements of an Id2a-retinoblastoma network in reprogrammed Müller glia that controls the expression of cell cycle genes and is required for transition from G1 to S phases of the cell cycle. In mutants, Müller glia that fail to divide undergo reactive gliosis, a pathological hallmark of Müller glia in mammals. Finally, we show that activation of the Midkine-a receptor, ALK, is required for Müller glia proliferation. These data provide mechanistic insights into Müller glia stem cells in the vertebrate retina and suggest avenues for eliciting neuronal regeneration in mammals.

## Introduction

Cell division is an essential biological process during tissue development, homeostasis and recovery from injury. In the central nervous system of adult mammals, stem cells reside in specialized niches, and whilst these cells maintain the ability to proliferate and generate new neurons they are unable to repair injuries(Kriegstein and Alvarez-Buylla, 2009; Ming and Song, 2011). In the vertebrate retina, Müller glia are among the last cells produced and they harbor molecular features of CNS stem and progenitor cells (Dyer and Cepko, 2000). In mammals, Müller glia respond to neuronal injury and death by undergoing partial dedifferentiation and entering the G1 phase of the cell cycle (Bringmann et al., 2006). However, this cellular reprogramming does not lead to cell division, and structural remodeling by Müller glia and the loss of retinal homeostasis are the typical sequalae (Bringmann et al., 2009; Hamon et al., 2016; Karl and Reh, 2010). Although pharmacological treatment or ectopic gene expression can elicit a modest neurogenic response in mammalian Müller glia, overall the regenerative capacity of these cells is very limited(Jorstad et al., 2017a; Ueki et al., 2015; Yao et al., 2018a). Importantly, in the limited instances where neuronal regeneration does occur, new neurons functionally integrate into existing synaptic circuits(Jorstad et al., 2017b; Yao et al., 2018b), indicating that in the mammalian retina the limitations of neuronal regeneration hinge on a more complete neurogenic response in Müller glia and Müller glia-derived neuronal progenitors.

In zebrafish, Müller glia function as intrinsic stem cells(Goldman, 2014; Gorsuch and Hyde, 2014; Hamon et al., 2016; Karl and Reh, 2010; Lenkowski and Raymond, 2014). In uninjured retinas, Müller glia reside in a mitotically quiescent state and function to maintain retinal homeostasis. Neuronal death triggers Müller glia to reprogram into a stem cell-like state, enter the cell cycle, and undergo a single asymmetric division to produce rapidly dividing, multipotent retinal progenitors with the ability to regenerate all types of retinal neurons (Goldman, 2014; Gorsuch and Hyde, 2014; Lenkowski and Raymond, 2014; Nagashima et al., 2013). Several signaling pathways have been identified that regulate the initial response of Müller glia(Goldman, 2014; Gorsuch and Hyde, 2014; Hamon et al., 2016; Karl and Reh, 2010; Lenkowski and Raymond, 2014). From this work, Ascl1, Lin28, and Stat3 have been identified as “core” transcriptional regulators that govern signaling cascades, including cytokine receptor signaling(Fausett and Goldman, 2006; Nelson et al., 2012; Ramachandran et al., 2010).

Midkine is a heparin-binding growth factor/cytokine that has multiple roles in neural development, tissue repair, and disease(Sakamoto and Kadomatsu, 2012; Sorrelle et al., 2017; Winkler and Yao, 2014). In several forms of cancers, Midkine promotes both proliferation and metastasis(Muramatsu, 2011). Midkine is also involved in inflammatory diseases by modulating immune cell activities(Herradon et al., 2019; Muramatsu, 2011; Weckbach et al., 2011). The diverse functions of Midkine are transduced through cell surface receptors, which may function as an individual trans-membrane protein or as members of a multi-protein complex(Muramatsu, 2011; Weckbach et al., 2011; Xu et al., 2014). During retinal development in zebrafish, *midkine-a*, one of two paralogous *midkine* genes in zebrafish, is expressed by retinal progenitors and functions as an upstream regulator of *id2a*, governing elements of the cell cycle(Calinescu et al., 2009a; Luo et al., 2012; Uribe and Gross, 2010). Post-mitotic neurons rapidly downregulate the expression of *midkine-a*. In adult zebrafish, retinal injury rapidly induces the expression of *midkine-a* in Müller glia and Müller glia-derived progenitors(Calinescu et al., 2009a; Gramage et al., 2014, 2015). Induction of *midkine-a* expression following injury has been reported for a variety of zebrafish tissues with the capacity to regenerate(Lien et al., 2006; Ochiai et al., 2004), suggesting that Midkine may universally regulate aspects of tissue regeneration. The molecular mechanisms whereby Midkine governs tissue regeneration are not well understood.

Using a Midkine-a loss-of-function mutant, we performed an initial transcriptome analysis, which revealed significant changes in pathways related to regulation of the cell cycle. Pursuing this *in vivo* demonstrated that in Müller glia Midkine-a is required during the single asymmetric division, as these cells progress from G1 to S phases of the cell cycle. Following the death of photoreceptors, Müller glia in Midkine-a mutants retain the ability to reprogram into a stem cell state and enter G1 phase of the cell cycle. However, analysis of the S phase cyclin, *ccn1a*, the cyclin dependent kinases, *cdk4* and *cdk6* and BrdU incorporation show that Midkine-a is required for Müller glia to progress beyond G1. Further, Midkine-a is required for upregulated expression of *id2a*, which inhibits the retinoblastoma (Rb) family of cell cycle inhibitors as Müller glia enter S-phase. The inability of zebrafish Müller glia to progress through the cell cycle results in selective failure in the regeneration of cone photoreceptors. In addition, the G1-arrested Müller glia undergo reactive gliosis and abnormal structural remodeling, hallmarks of Müller glia pathology in the mammalian retina. Finally, we provide evidence that activation of the the Midkine receptor, Anaplastic Lymphoma Kinase (Alk), is required for proliferation in Müller glia.

## Results

### Loss-of function mutant, *mdka*^*mi5001*^

We generated a CRISPR-Cas9-mediated Midkine-a loss-of-function mutant, *mdka*^*mi5001*^, which carries a 19 bp deletion in the exon three of *midkine-a*. This deletion results in a predicted premature stop codon (Fig. S1A and B) and absence of protein in Western blot analysis (Fig. 1A). and larval and adult retinas (Fig. S1C). *Mdka*^*mi5001*^ larvae progress normally through early developmental stages and at 48 hpf show only slight reduction in body pigmentation, shortened body length, and smaller eyes (Fig. 1B and S2A and B). The pigmentation defect recovers by 72 hpf (Fig. 1B). Notably, the *mdka*^*mi5001*^ mutants replicate the delayed retinal development described previously following morpholino-mediated knock-down of Midkine-a (Fig. 1C;(Luo et al., 2012)).

**Figure 1.**
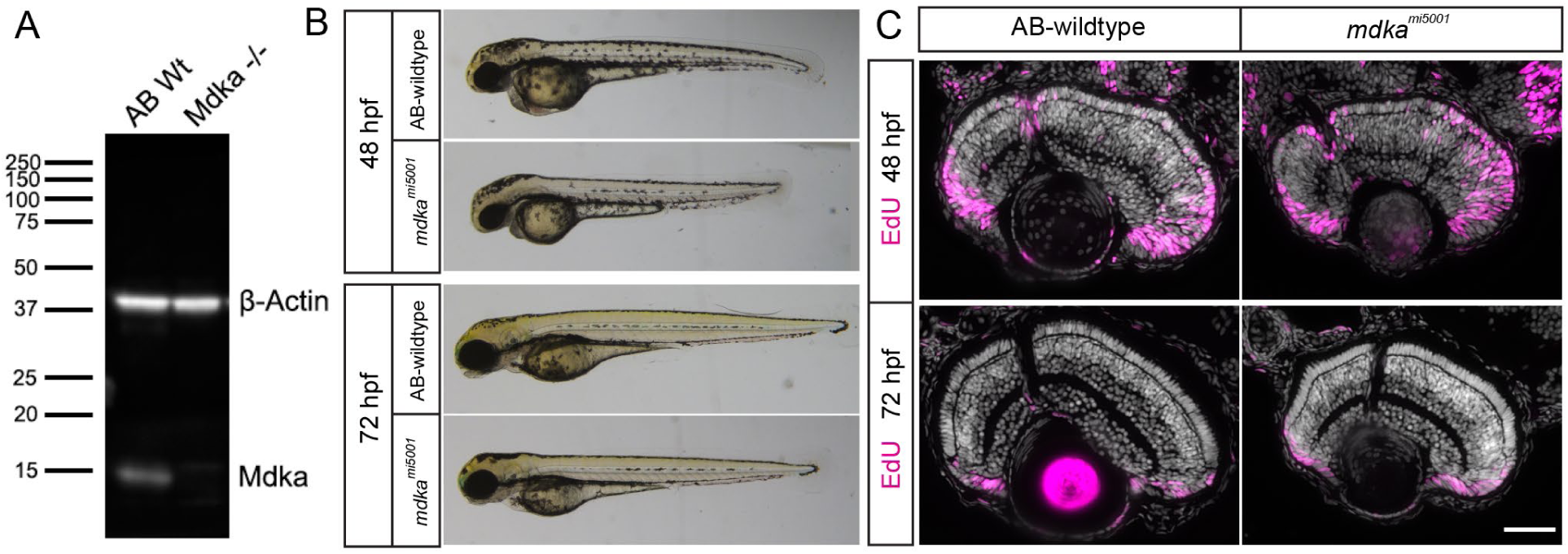
Midkine-a governs cell cycle kinetics during retinal development. (A) Using larvae at 48 hours post fertilization (hpf), Western blot analysis for Midkine-a confirms lack of protein in the *mdka*^*mi5001*^ mutant. (B) Lateral views of larvae at 48 and 72 hpf: AB-wildtype and *mdka*^*mi5001*^ mutant. (C) Proliferation assay with EdU labeling at 48 hpf. Compared to wildtype, there is an increased number of EdU-labeled cells in the retinas of *mdka*^*mi5001*^ mutants. Retinal lamination and cellular differentiation are delayed, but not not blocked in the *mdka*^*mi5001*^ at 72 hpf. Scale bar; 40 μm. See also Figure S1 and S2.

### Transcriptome Analysis

Larvae from adult *mdka*^*mi5001*^ mutants were initially evaluated using transcriptome analysis of whole embryos at 30 hpf. This identified 638 differentially expressed genes (log2 fold change ≥1 and a false discovery rate ≤0.05) (Table S1). Pathway level analysis with the Reactome tool identified 181 pathways that were differentially regulated (Table S2). Of the 157 differentially-regulated pathways, 33 were related to cell cycle regulation (Table S2). These data validated our previous study (Luo et al., 2012) and directed us to evaluate the role of Midkine-a in regulating proliferation in Müller glia.

### Regeneration of cone photoreceptors is severely compromised in the *mdka*^*mi5001*^ mutant

Persistent, growth-associated neurogenesis is a hallmark of teleost fish(Hitchcock et al., 2004). In the growing eye and retina, stem and progenitor cells at the ciliary marginal zone generate new retinal neurons, with the exception of rod photoreceptors(Cerveny et al., 2012). Fate-restricted, proliferating rod precursors, sequestered in the outer nuclear layer, selectively give rise new rod photoreceptors(Raymond and Rivlin, 1987; Stenkamp, 2011). This growth-associated neurogenesis occurs normally in the retinas of *mdka*^*mi5001*^ mutants, and there is no apparent alteration in the maturation or variety of cell types in the *mdka*^*mi5001*^ retina (Fig. 1C). In response to neuronal cell death, Müller glia in zebrafish dedifferentiate and undergo a single asymmetric division to produce retinal progenitors, which rapidly divide, migrate to areas of cell loss and differentiate to replace the ablated neurons(Nagashima et al., 2013). To assess photoreceptor regeneration in the *mdka*^*mi5001*^, we used a photolytic lesion that selectively kills photoreceptors; photoreceptors undergo apoptotic cell death by 1 day post-lesion (dpl)(Bernardos et al., 2007; Vihtelic and Hyde, 2000). In wildtype retinas, by 1 dpl Müller glia can be labeled with antibodies against the proliferation marker, PCNA, and by 3 dpl the Müller glia-derived progenitors form radial neurogenic clusters that span the inner nuclear layer (Fig. 2A). In contrast, in the *mdka*^*mi5001*^ retinas, PCNA labeling was completely absent at 1 dpl, and only a few cells were PCNA-positive at 3 dpl (Fig. 2A and Fig. S3A-B). However, at 5 dpl there was no difference in the number of PCNA+ cells in the outer nuclear layer of wildtype and mutant retinas (Fig. S3B and C). In wildtype retinas, the PCNA+ cells in the outer nuclear layer are a mixture of Müller glia-derived cone progenitors and endogenous, proliferating rod precursors. We infer tha in mutants, the dividing cells in the outer nuclear layer at 5dpl consist solely of rod precursors (see below).

**Figure 2.**
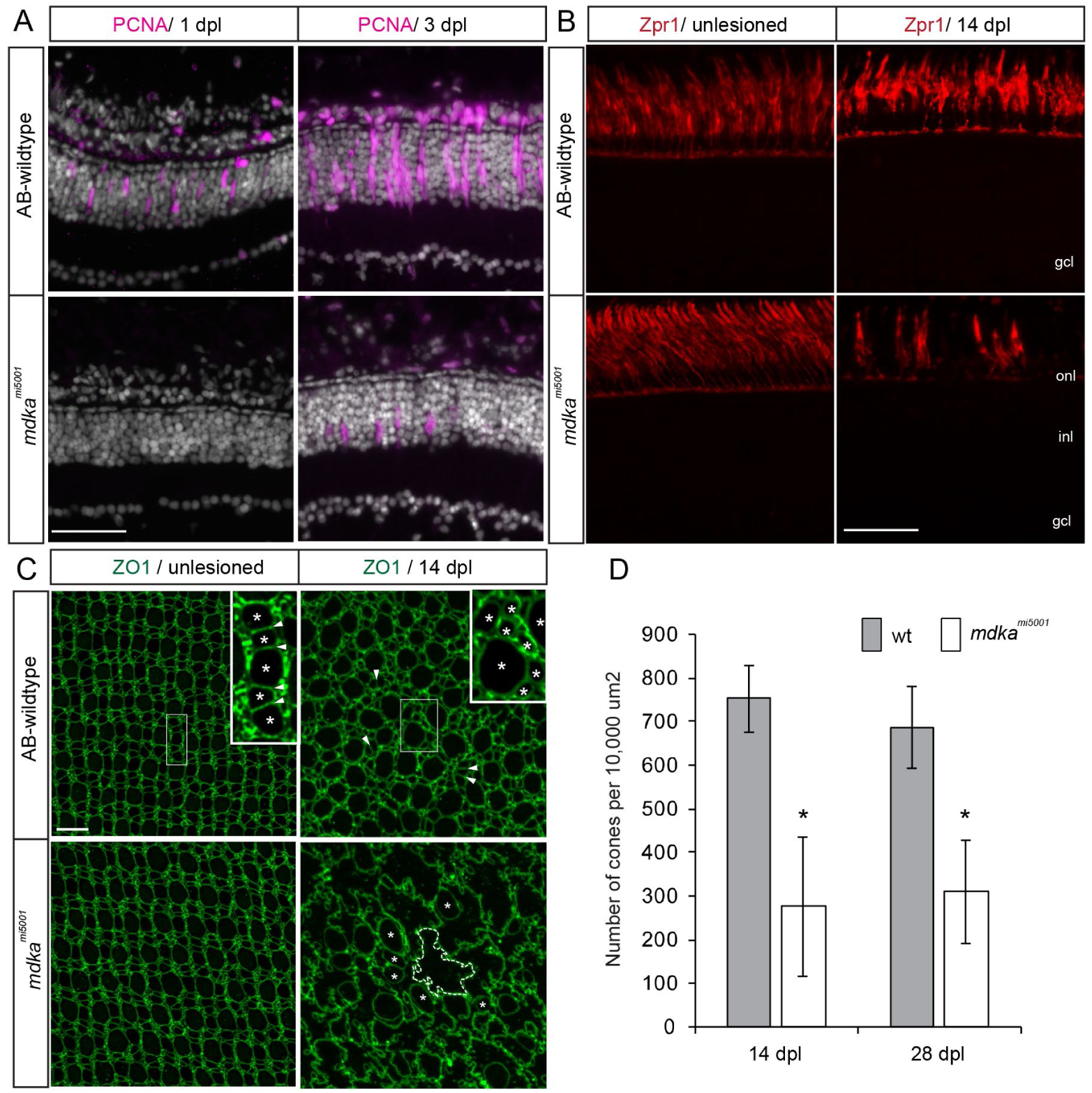
In the *mdka*^*mi5001*^ mutant, Müller glia fail to proliferate and cone regeneration is compromised. (A) Immunocytochemistry for PCNA (magenta) in wildtype and *mdka*^*mi5001*^ at 1 and 3 days post lesion (dpl). In wildtype, Müller glia become positive for PCNA at 1 dpl. At 3 dpl, PCNA+ progenitors form neurogenic clusters. Mutant retinas lack PCNA-labeled cells at 1 dpl. Very few, isolated cells are positive for PCNA in the *mdka*^*mi5001*^ at 3 dpl. (B) Immunocytochemistry for red and green cone marker Zpr1 (red) in wildtype and *mdka*^*mi5001*^ retinas. At 14 dpl very few Zpr1+ photoreceptors have regenerated in the *mdka*^*mi5001*^ mutant compared with wildtype. (C) Flat-mounted retinal preparation immunostained with ZO1 in unlesioned and 14 dpl. In unlesioned retina of both wildtype and *mdka*^*mi5001*^, cone photoreceptors form a crystalline mosaic array in the planar apical surface of the retina(Livak and Schmittgen, 2001; Nagashima et al., 2017). Higher magnification of boxed region indicates the alignment of cones in the mosaic array (asterisks) with flattened cell boundaries (arrowheads). At 14 dpl in wildtype, cone photoreceptors regenerate (asterisks), although the crystalline mosaic array is not restored. In the *mdka*^*mi5001*^ retina cone profiles are instead replaced by irregularly shaped, expanded Müller glial apical processes (dotted line). (D) Counts of ZO1-labeled cone photoreceptors at 14 and 28 dpl. Significantly fewer cones are regenerated in the *mdka*^*mi5005*^ mutant (white) compared with wildtype (gray). *p<0.0001. onl: outer nuclear layer; inl: inner nuclear layer; gcl: ganglion cell layer. Scale bars; 30 μm (A, B); 10 μm (C). See also Figure S3 and S4.

We next assayed photoreceptor regeneration using specific cone and rod photoreceptor markers, Zpr1 and Zpr3, respectively, and flat-mount retinal preparations immunostained with the cell junction marker, ZO1 (Zonula Occuludens 1). In wildtype retinas, regenerated cones appear as early as 5 dpl, and by 14 dpl the regeneration of cone photoreceptors is largely complete (Fig. 2B and Fig. S4A). In contrast, at 5 dpl in the *mdka*^*mi5001*^ retinas, regenerated cones are nearly completely absent, and at 7 dpl only a few immature cone photoreceptors are present (Fig. S4A). The relative lack of cones persists through 14 and 28 dpl, demonstrating that in the mutant retinas regeneration of cone photoreceptors is blocked, not simply delayed (Fig. 2D and S4A). These results indicate that cone photoreceptor regeneration is permanently compromised in the absence of Midkine-a. In contrast, rod photoreceptors are regenerated slowly but to apparently normal numbers in the *mdka*^*mi5001*^ mutants (Fig. S4B). In wildtype retinas, mature rod photoreceptors appear by 7 to 14 dpl (Fig. S4B), whereas in the *mdka*^*mi5001*^ mutants, rod photoreceptors are immature at 7dpl, but completely replenished by 28 dpl (Fig. S4B). Taken together, these results demonstrate that Midkine-a is required for Müller glial stem cells to proliferate, and the absence of Midkine-a leads to a failure of cone photoreceptor regeneration. In contrast, rod photoreceptors regenerate normally in the mutants, and we infer they arise from the fate-restricted, proliferating rod precursors that persist in the depleted outer nuclear layer.

### Müller glia in the *mdka*^*mi5001*^ mutants undergo gliotic remodeling

In all vertebrate retinas, neuronal death induces a gliotic response in Müller glia(Bringmann et al., 2006, 2009; Jadhav et al., 2009). Although the initial reactive gliosis is neuroprotective, persistent gliosis results in dysregulation of retinal homeostasis, glial remodeling and scar formation, and the subsequent death of neurons(Bringmann et al., 2006). In zebrafish, the gliotic response of Müller glia is transient and interrupted by cell cycle re-entry(Thomas et al., 2016). To determine if the failure of Müller glial to enter the cell cycle in Midkine a mutants leads to a mammalian-like gliotic response, the expression of glial fibrillary acidic protein (GFAP) was compared in the wildtype and *mdka*^*mi5001*^ retinas. In unlesioned retinas, immunostaining for GFAP labels the basal processes of Müller glia(Bernardos and Raymond, 2006). In wildtype retinas at 28 dpl, GFAP immunolabeling resembles that in unlesioned retinas. In contrast, in the *mdka*^*mi5001*^ retinas at 28 dpl, GFAP immunolabeling is present throughout the cytoplasm, extending apically into the inner nuclear layer (Fig. S5A). Enhanced expression of GFAP is a marker of gliosis in mammalian Müller glia(Bringmann et al., 2006). To further characterize the gliotic response in the mutant retinas, the transgenic reporter, *Tg(gfap:eGFP)*^*mi2002*^, was crossed into the *mdka*^*mi5001*^ background(Bernardos and Raymond, 2006). Computing the planimetric density of Müller glia in flat-mount preparations at 28 dpl showed no significant difference in the number of Müller glia in control and mutant retinas, as expected. However, in the *mdka*^*mi5001*^*/Tg(gfap:eGFP)*^*mi2002*^ line Müller glia remained hypertrophic, as evidenced by elevated e*GFP* levels (Fig. S5B, arrows, and Supplementary Movie 1), and these cells adopt abnormal morphologies, including expanded lateral extensions in the inner plexiform layer (Fig. S5C) and migration of the somata into the outer plexiform layer (Fig. 3A,B and Movie 2, 3). This hypertrophic morphology is also revealed in flat mounts stained with the cell junction marker, ZO1, in which the apical profiles of Müller glia have expanded to fill the planar surface of the outer limiting membrane previously occupied by cones (Fig. 2C, lower left panel). The abnormal gliotic remodeling observed in the mutant retinas is a hallmark of persistent reactive gliosis in mammals.

**Figure 3.**
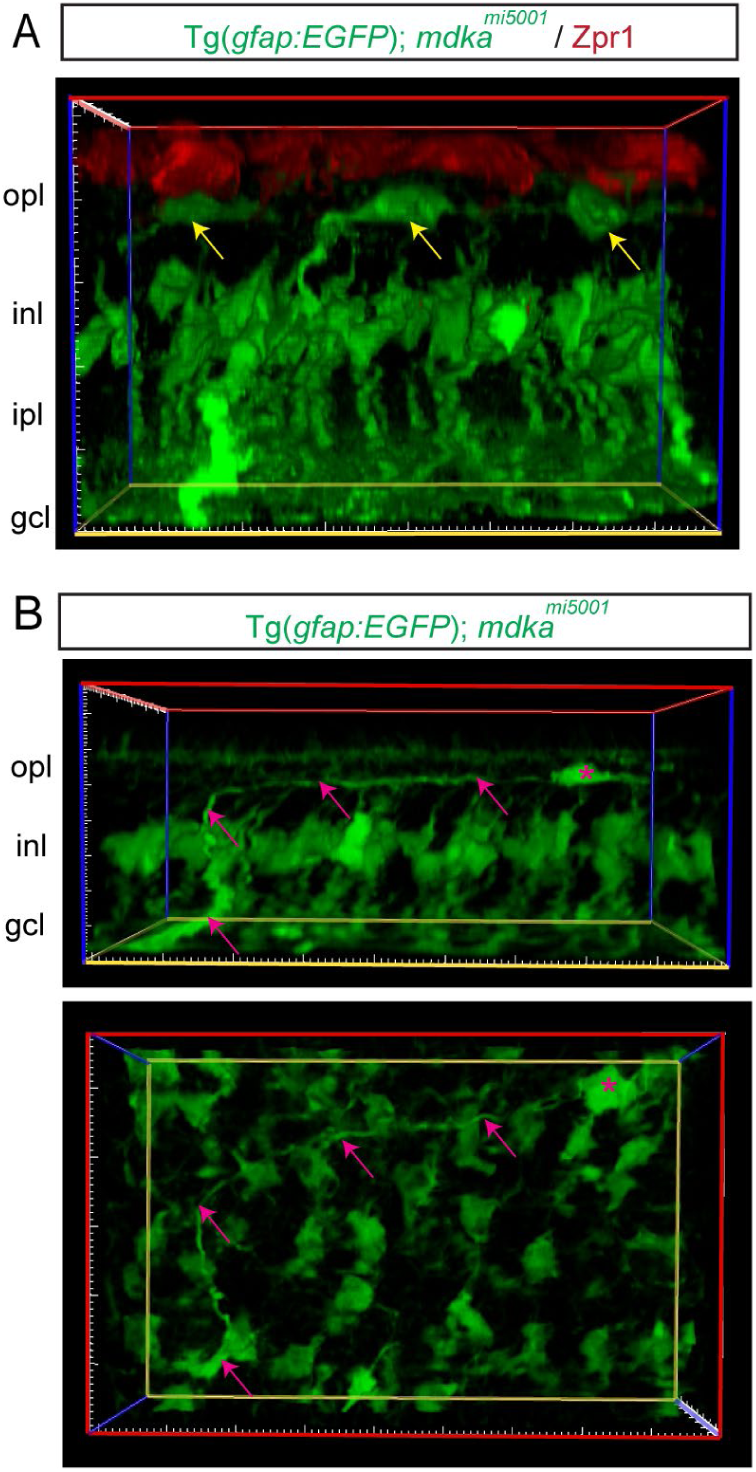
Following photoreceptor death, Müller glia in the *mdka*^*mi5001*^ mutant undergo gliotic remodeling. (A) Cross section view of 3D reconstructed image in the Tg(*gfap:GFP*); *mdka*^*mi5001*^ (green) retina at 28 days post lesion (dpl), immunolabeled with Zpr1 (red) in a flat-mount preparation. Yellow arrows indicate displaced Müller glia somata in the outer plexiform layer. (B) Cross section and flat-mounted views of the 3D reconstructed image. Displaced Müller glia (magenta asterisk) retain basal radial process (magenta arrows). opl: outer plexiform layer; inl: inner nuclear layer; ipl: inner plexiform layer; gcl: ganglion cell layer. See also Figure S5.

### Müller glia in the the *mdka*^*mi5001*^ dedifferentiate in response to photoreceptor death

In response to neuronal cell death, Müller glia spontaneously reprogram, upregulating stem cell-associated genes prior to entering the cell cycle(Goldman, 2014; Gorsuch and Hyde, 2014; Hamon et al., 2016; Lenkowski and Raymond, 2014). Immunostaining retinas at 1 and 2 dpl for the stem-cell associated proteins, Rx1 and Sox2, labeled elongated, polygonal nuclei, characteristic of Müller glia(Nagashima et al., 2013)(Gorsuch et al., 2017), in both wildtype and mutant retinas (Fig. 4A). Further, the Rx1 and Sox2 positive nuclei were displaced apically in both (Fig. 4A and Fig. S6), revealing the interkinetic nuclear migration that is associated with cell cycle progression in Müller glia(Nagashima et al., 2013). We also evaluated the reprogramming in Müller glia by qPCR for the core transcriptional factors, *ascl1a, lin28 and stat3 (Fausett and Goldman, 2006; Nelson et al., 2012; Ramachandran et al., 2010)*. At 30 and 36 hpl, which is prior to when Müller glia divide, *ascl1a, stat3* and *lin28* are significantly upregulated in both wildtype and mutant retinas, although the expression level of *ascl1a* is slightly reduced in the mutants (Fig. 4B-D). These data indicate that in the absence of Midkine-a Müller glia respond to photoreceptor death by reprogramming into a stem cell-like state.

**Figure 4.**
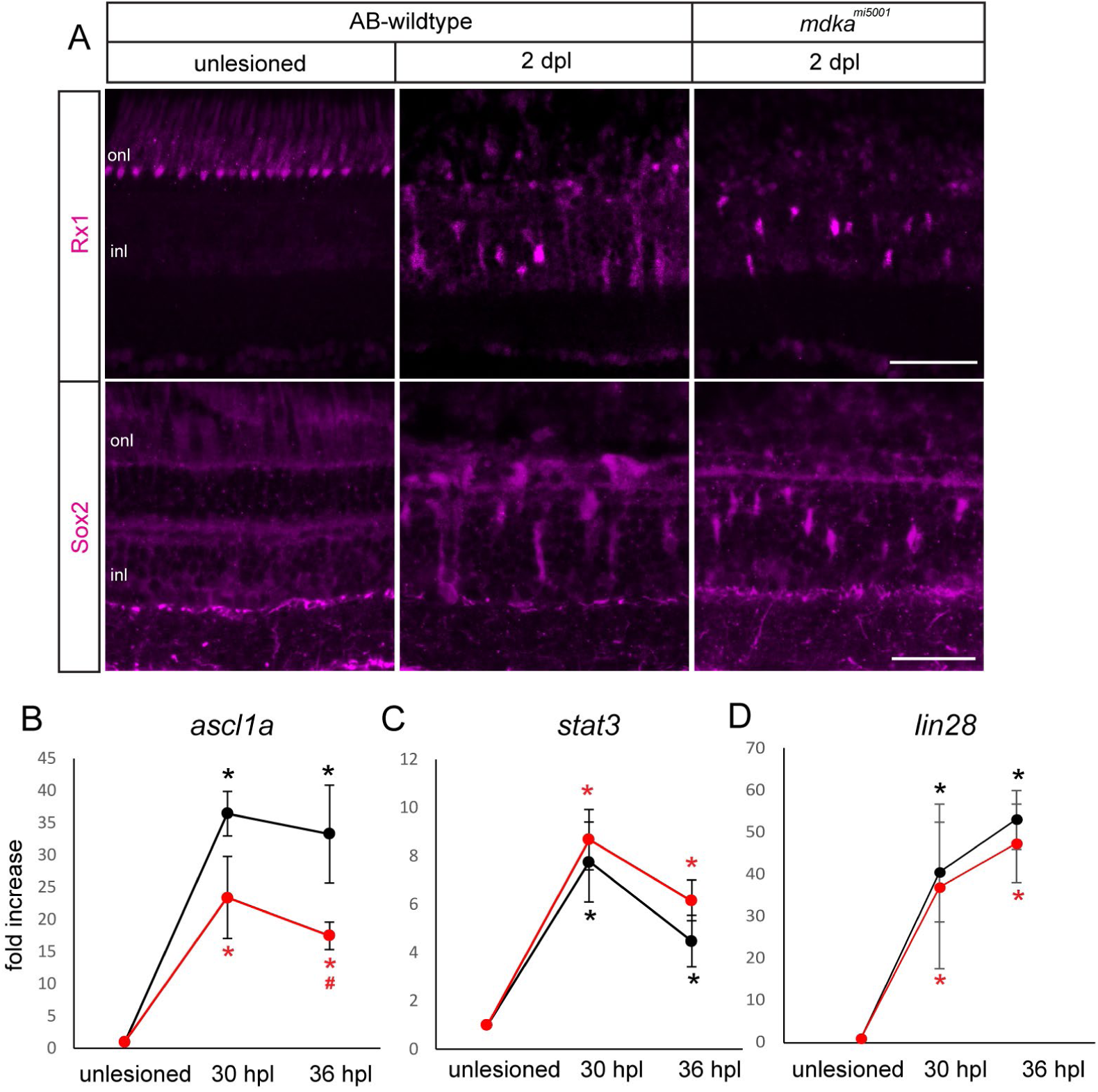
Müller glia in the *mdka*^*mi5001*^ mutant dedifferentiate following photoreceptor death. (A) Immunocytochemistry for regeneration-associated genes, Rx1 and Sox2, following photolytic lesion in wildtype and *mdka*^*mi5001*^ retinas. Lesion induces Rx1 and Sox2 expression in Müller glia both in wildtype and *mdka*^*mi5001*^ retinas at 2 dpl. (B-D) qPCR for dedifferentiation markers, *ascl1a, stat3*, and *lin28*, at 30 and 36 hpl. Both wildtype and *mdka*^*mi5001*^ upregulate *ascl1a* (B), *stat3* (C), and *lin28* (D) following lesion. *p<0.03 relative to unlesioned. ^#^p=0.0066 relative to wildtype. Scale bars; 30 μm (A). onl: outer nuclear layer; inl: inner nuclear layer. See also Fig. S7.

### Midkine-a is partially responsible for *ascl1a* expression via phosphorylation of Stat3

Following a retinal lesion, phosphorylation of Stat3 is required for the upregulation of *ascl1a* in Müller glia(Nelson et al., 2012; Zhao et al., 2014). Consistent with previously published data (Kassen et al., 2007; Zhao et al., 2014), at 1 and 2 dpl, pStat3 antibodies label Müller glia in wildtype animals (Fig. S7). In contrast, in *mdka*^*mi5001*^ retinas at 1 and 2 dpl, pStat3 immunostaining was markedly diminished (Fig. S7), demonstrating that in Müller glia Midkine-a acts upstream of pStat3, which can explain the reduced expression of *ascl1a* in the mutant retinas.

### Cell cycle arrest in mutant Müller glia

Following reprogramming, Müller glia begin entering the cell cycle at 24 hpl and complete the asymmetric cell divisions by 42 hpl (Nagashima et al., 2013). We next asked whether Müller glia in the *mdka*^*mi5001*^ possess the ability to enter the cell cycle by quantifying the expression of G1 phase cyclins, *cyclin d1(ccnd1)* and *cyclin e1 (ccne1)*. These cyclins are expressed during G1 and function to drive G1 to S phase transition(Dyer and Cepko, 2001). We isolated mRNA at 30 and 36 hpl, knowing that cell cycle progression is not completely synchronous among the population of Müller glia, but that these time points will allow us to capture gene expression changes in Müller glia and exclude Müller glia-derived progenitors. This analysis showed that *mdka*^*mi5001*^ upregulates *ccnd1* and *ccne1* significantly at both 30 and 36 hpl (Fig. 5A and 5B), indicating that following photoreceptor death in the *mdka*^*mi5001*^ mutants Müller glia enter the G1 phase of the cell cycle. In wildtype retinas, cell cycle entry is followed by upregulation of S phase cyclin, *ccna2* (Fig. 5C). In contrast, there is no upregulation of *ccna2 in the mdka*^*mi5001*^ retinas, indicating that Müller glia in mutants fail to progress from G1 to S (Fig. 5C). This was confirmed using the S-phase label, BrdU, between 24 to 30 hpl. In wildtype retinas, Müller glia are uniformly labeled with BrdU, whereas in the in the *mdka*^*mi5001*^ retinas there are no BrdU-labeled cells (Fig. 5D, n=6 retinas). Consistent with these results, the expression of the cell cycle regulators, *cyclin-dependent kinase 4* and *6*, are dysregulated in mutant retinas (Fig. 5E and 5F). Together, these results indicate that following photoreceptor cell death in the *mdka*^*mi5001*^ retinas, Müller glia undergo cell cycle arrest in the G1 phase, demonstrating that in reprogrammed Müller glia Midkine-a is required for the G1-S phase transition.

During retinal development, Midkine-a governs cell cycle kinetics through Id2a(Luo et al., 2012). Id proteins play important roles in cell cycle regulation during development and in cancer(Lasorella et al., 1996; Sikder et al., 2003). In wildtype retinas, *id2a* expression is markedly upregulated at 30 hpl, as Müller glia progress through the cell cycle, and rapidly returns to baseline levels by 48 hpl, when the single asymmetric mitotic division is complete (Fig. 5G). This transient induction of *id2a* is completely absent in the *mdka*^*mi5001*^ retinas (Fig. 5G). In cancer cells, Id2 proteins antagonize the retinoblastoma (Rb) family of cell cycle inhibitors, thereby allowing progression from G1 to S phase of the cell cycle. (Lasorella et al., 2001; Sikder et al., 2003). Previous analyses of the Müller glia specific transcriptome show that *p130*, one of the Rb gene family, exhibits highest expression among Rb genes in quiescent Müller glia(Nieto-Arellano and Sánchez-Iranzo, 2018; Sifuentes et al., 2016). Consistent with these data, we validated that in wildtype retinas the expression of p130 decreases as Müller glia progress through the cell cycle (Fig. 5H). In contrast, in *mdka*^*mi5001*^ retinas at 30 and 36 hpl, p130 levels are elevated above the those found in quiescent Müller glia (Fig. 5H). These results suggest that Id2a is downstream of Midkine-a, and in Müller glia Id2a functions to inhibit retinoblastoma genes.

**Figure 5.**
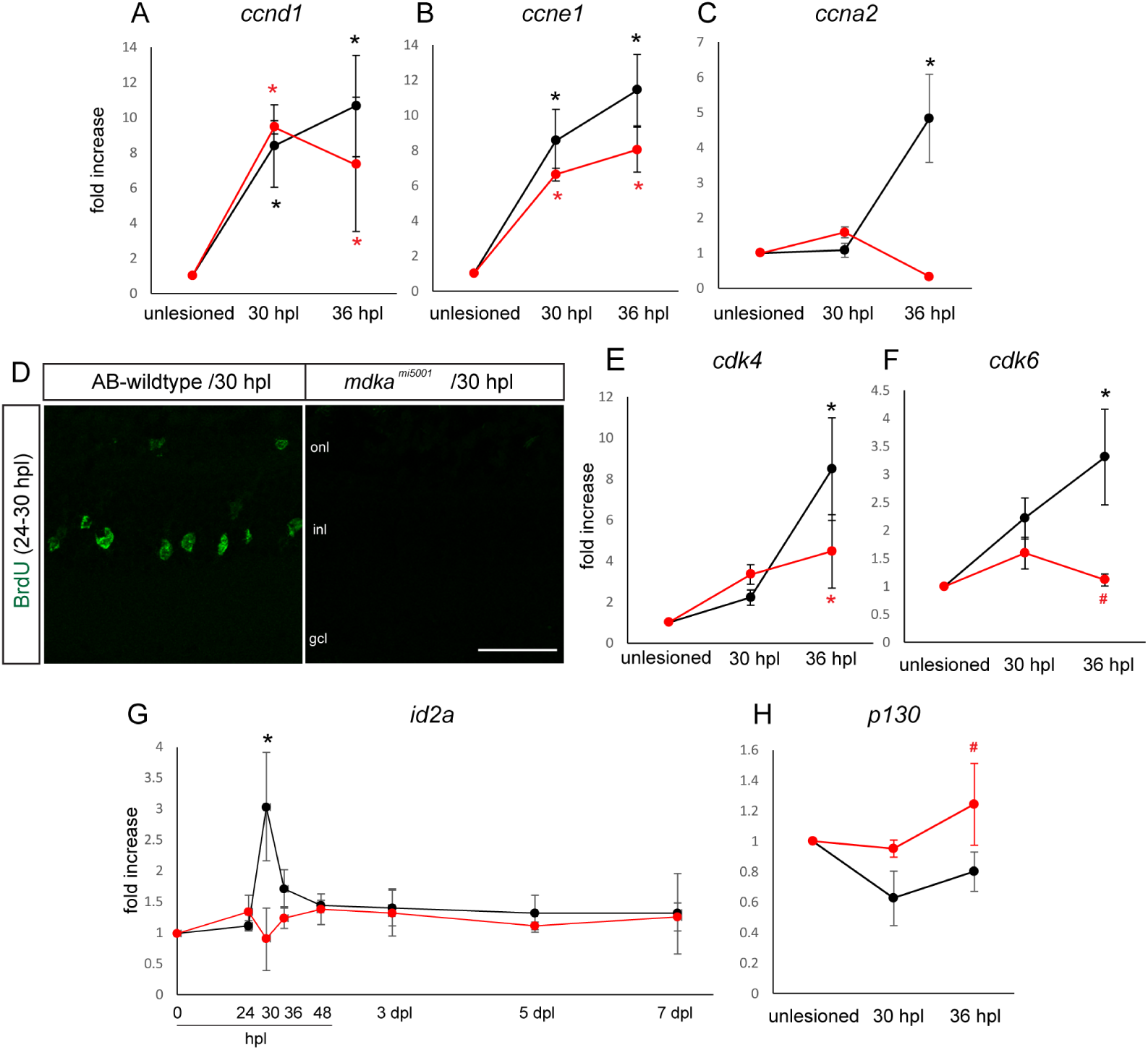
Müller glia in the *mdka*^*mi5001*^ mutant arrest in the G1 phase of the cell cycle. (A-C) qPCR assay for cell cycle regulator cyclins, *ccnd1, ccne1*, and *ccna2* following photolytic lesion. Both wildtype and *mdka*^*mi5001*^ upregulate G1 cyclins, *ccnd1* and *ccne1*, following lesion (A-B). The *mdka*^*mi5001*^ mutant fails to upregulate S phase cyclin, *ccna2* (C). S phase assay with BrdU labeling (green) between 24 to 30 hours post lesion. Müller glia in the *mdka*^*mi5001*^ mutants did not incorporate BrdU following lesion. (E-H) qPCR of additional cell cycle regulators. Expression of cyclin dependent kinases, *cdk4* and *cdk6*, are dysregulated in the *mdka*^*mi5001*^ retinas (E-F). The wildtype transiently upregulates *id2a* at 30 hpl, whereas expression levels did not change in the *mdka*^*mi5001*^ (G). Expression of the cell cycle inhibitor, *p130*, decreases in the wildtype after lesion, while *mdka*^*mi5001*^ maintains steady levels of expression (H). *p<0.03 relative to unlesioned. *p<0.03 relative to wildtype. Scale bars; 30 μm (D).

### Anaplastic lymphoma kinase is responsible for Müller glial proliferation

Anaplastic Lymphoma Kinase (ALK) is a superfamily of receptor tyrosine kinases, originally identified as an oncogene in human lymphomas. ALK is involved in the initiation and progression of many cancers, including neuroblastoma(Hallberg and Palmer, 2013; Morris et al., 1995; Webb et al., 2009). Midkine, and its related protein Pleiotrophin, are the only ligands known to activate ALK(Stoica et al., 2001, 2002). To determine if Alk functions as a Midkine-a receptor on Müller glia during photoreceptor regeneration, double immunocytochemistry was performed for phosphorylated Alk (pAlk) and PCNA following a photolytic lesion. In wildtype retinas, pAlk colocalizes with PCNA, indicating activation of Alk in dividing Müller glia and Müller glia-derived progenitors (Fig. 6A). In contrast, both pAlk and PCNA immunolabeling were absent in *mdka*^*mi5001*^ retinas, indicating that in the retina Midkine-a is required for ALK phosphorylation. To test if activation of ALK is required for proliferation among Müller glia, wildtype animals were housed from 24 to 72 hpl in the ALK inhibitor, TAE684. Inhibiting the activation of ALK phenocopied the proliferation defect observed in the *mdka*^*mi5001*^ mutants (Fig. 6B and 6C). These data indicate that phosphorylation of Alk is required for Müller glia to proliferate and identify ALK as a putative receptor for Midkine-a during the initial asymmetric division in Müller glia and the subsequent regeneration of cone photoreceptors.

**Figure 6.**
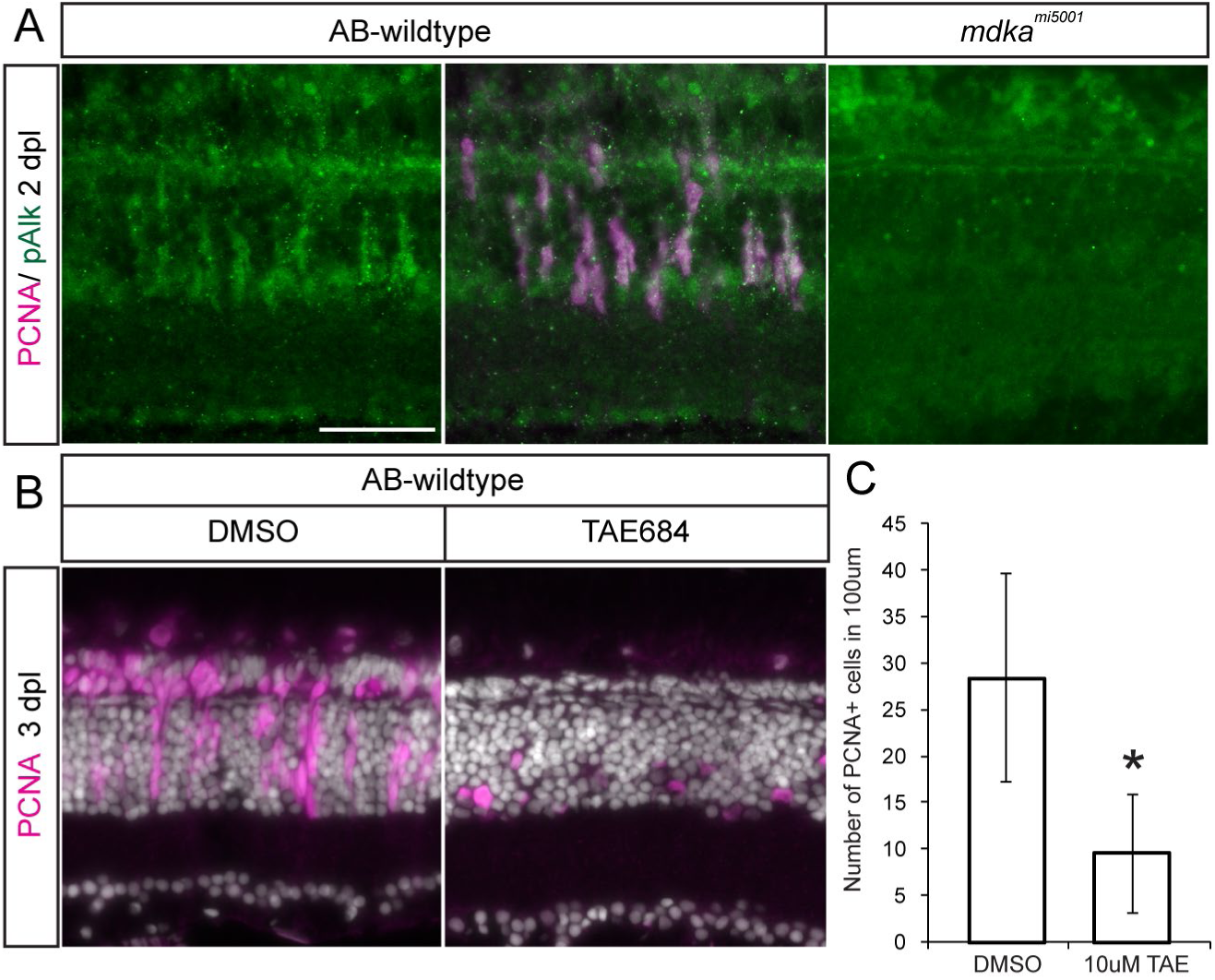
Activation of the Anaplastic Lymphoma Kinase receptor is required for Müller glial to proliferate. (A) Double-immunostaining for PCNA (magenta) and phosphorylated-Anaplastic lymphoma kinase (pAlk, green). PCNA+ cells express pAlk in wildtype retina at 2 dpl, whereas pAlk immunostaining was not detected in *mdka*^*mi5001*^ retina. (B) Pharmacological inhibition of Alk using TAE684 suppresses proliferation in the wildtype retinas. (C) Counts of PCNA-positive proliferative cells in DMSO or TAE684 treated wildtype retinas at 3 dpl. t-test, *p=0.0033. Scale bars; 30 μm (A).

## Discussion

In mammals, neuronal damage in the retina stimulates transient entry of Müller glia into the cell cycle, however, any subsequent proliferative response is very limited (Dyer and Cepko 2000; Hamon et al. 2019; Rueda et al. 2019). In zebrafish, Müller glia respond to neuronal death by spontaneously entering and transiting the cell cycle, giving rise to Müller glia-derived progenitors that amplify in number and functionally replace ablated neurons. Numerous studies have identified transcriptional regulators and signaling cascades that promote Müller glia reprogramming in both mammals and fish(Goldman, 2014; Gorsuch and Hyde, 2014; Hamon et al., 2016; Karl and Reh, 2010; Lenkowski and Raymond, 2014). The molecular mechanisms that govern cell cycle kinetics in Müller glia, while essential, have received relatively little attention. Here we provide the first evidence that Midkine-a, ostensibly acting as an autocrine regulator of proliferation, is required in Müller glia for the transition from G1 to S phases of the cell cycle.

Our data support the mechanistic model shown in Fig. 7. In unlesioned retinas, Müller glia remain quiescent in the G0 phase (Fig. 7A). In response to photoreceptor cell death, nearby Müller glia upregulate reprogramming-associated genes, and enter the cell cycle (Fig. 7B). Midkine-a, signaling through Alk receptors, promotes the expression of *ascl1a* via phosphorylation of Stat3 (Fig. 7B). Midkine-a also induces the brief, transient upregulation of Id2a, which suppresses cell cycle inhibitor, p130, thereby allowing Müller glia to enter both S and the subsequent phases of the cell cycle (Fig. 7B). In Midkine-a loss of function mutants, Müller glia initiate reprogramming into a stem cell state and enter G1 phase of the cell cycle, but fail to activate Id2a and fail to transition from G1 to S and mitosis (Fig. 7B, C). The consequence of this cell cycle block is the selective failure in the regeneration of cone photoreceptors.

**Figure 7.**
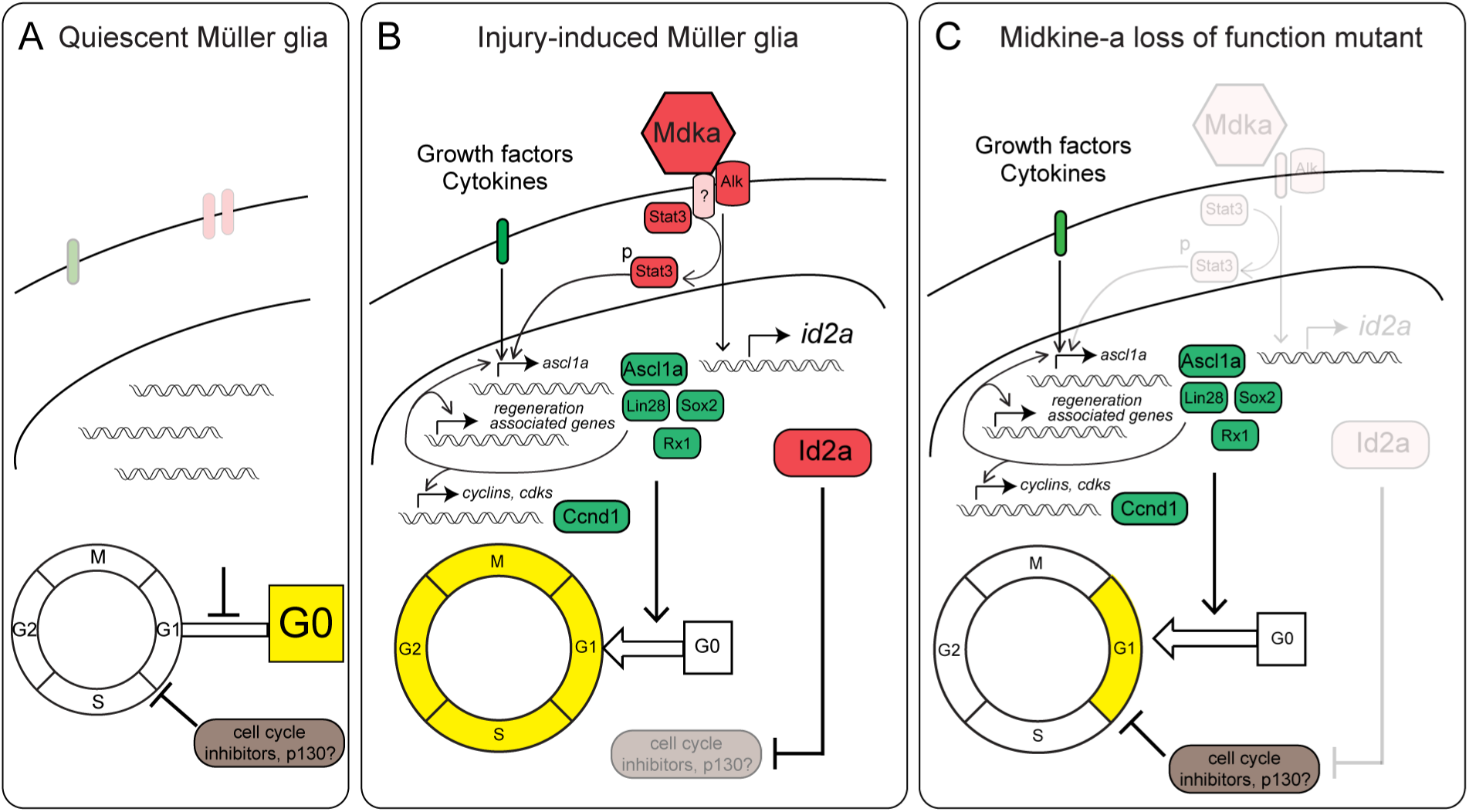
Model of Midkine-a-mediated cell cycle regulation in Müller glia. (A) In quiescent Müller glia, cell cycle inhibitors keep cells in G0 phase. (B) Injury induces cytokines and growth factors to upreuglate regeneration-associated reprogramming genes for dedifferentiation and cell cycle reentry. Midkine-a-Alk signaling participates in the induction of the regeneration-associated gene, *ascl1a*, via phosphorylation of Stat3. Midkine-a signaling also induces expression of the cell cycle regulator, *id2a*, that inhibits cell cycle inhibitors, such as p130. (C) In the absence of Midkine-a, Müller glia fail to suppress cell cycle inhibition, resulting in G1 arrest.

During retinal development, Müller glia emerge from late stage retinal progenitors, and cell cycle inhibitors play pivotal roles in their fate determination(Turner and Cepko, 1987; Furukawa et al., 2000; Levine et al., 2000; Ueki et al., 2012; Del Debbio et al., 2016). In adult retinas, Müller glia retain an intrinsic genetic program that is shared with these retinal progenitors(Roesch et al., 2008). Müller glia in mammalian retinas respond to neuronal death by entry into the cell cycle (Dyer and Cepko 2000; Hamon et al. 2019; Rueda et al. 2019; Joly et al., 2011; Nomura-Komoike et al., 2016). Importantly, in the mouse, genetic modifications that allow Müller glia to persistently express *cyclin* genes is sufficient to promote their mitotic division (Hamon et al., 2019; Rueda et al., 2019). Our data, together with these observations, suggest that Müller glia in both mammals and zebrafish posses similar mechanisms that function to integrate injury-related signals from dying neurons and promote entry into the cell cycle, but in zebrafish, Midkine-a serves as a unique autocrine signal that allows these cells to proliferate.

Following photoreceptor death, reprogrammed Müller glia enter the cell cycle within 24 hpl and undergo a single division by 42 hpl(Nagashima et al., 2013). This temporal sequence is closely coordinated, but not completely synchronized. We timed experiments using RNA isolated from whole retinas such that cell cycle-related gene expression could be evaluated in Müller glial stem cells, while excluding Müller glia-derived progenitors. Following photoreceptor death in zebrafish, *id2a* is upregulated transiently at 30 hpl and returns to baseline levels by 36 hpl. This indicates that Midkine-a-dependent Id2a expression is required for the proliferation of Müller glia, but not Müller glia-derived progenitors. The family of Id proteins is involved in intrinsic control of proliferation during development and in cancer(Ruzinova and Benezra, 2003; Sikder et al., 2003). Id2 regulates the Rb proteins, and high levels of Id2 can suppress the Rb tumor suppressor pathway, which blocks progression from G1 to S phase of the cell cycle(Lasorella et al., 2000, 2001). In adult mice, the Rb-family member p130 maintains quiescence in muscle satellite cells, which retain the capacity to self renew and regenerate myoblasts(Carnac et al., 2000). During sensory hair cell regeneration in zebrafish inner ear, p130 is downregulated immediately following injury(Jiang et al., 2014). Consistent with these data, our *in silico* screening identified p130 as a highly expressed gene in quiescent Müller glia, suggesting p130 expressed in Müller glia of uninjured retinas functions to restrict their proliferation(Nieto-Arellano and Sánchez-Iranzo, 2018; Sifuentes et al., 2016). Our data suggest that in response to cell death Midkine-a signaling blocks p130 through upregulating Id2a, allowing Müller glia to progress through the cell cycle. We suggest the brief up- and down-regulation of *id2a* is a mechanism that allows Müller glia to divide, but restricts these cells to a single mitotic cycle. Further, our data suggest Midkine-a is required for the rising phase of this transient *id2a* expression. Notably, increased levels of Id2 are also present in anaplastic large cell lymphomas that result from constitutive activation of the Midkine receptor, ALK(Mathas et al., 2009). It is not known if ALK in Müller glia functions alone or as a member of multi-protein complex to relay the Midkine-a signal. Receptor protein tyrosine phosphatase-zeta (RPTP-ζ) is also a known receptor for Midkine and can activate the intracellular kinase domain of ALK, and may function as a co-receptor to transduce Midkine-a signaling in Müller glia(Hallberg and Palmer, 2013; Mathas et al., 2009).

Following photoreceptor death in the *mdka*^*mi5001*^ mutants, reprogrammed Müller glia undergo cell cycle arrest, whereas proliferation among the fate-restricted rod precursors within the outer nuclear layer is unimpeded. As a result, the Müller glia-derived regeneration of cone photoreceptors fails, but rod photoreceptors are eventually replenished. This observation is consistent with previous reports that also identified separate lineages between regenerated cone and rod photoreceptors(Gorsuch et al., 2017; Morris et al., 2008; Thummel et al., 2010).

Heterogeneity of stem and progenitor populations is a clinical challenge when treating malignant tumors, where Midkine is highly expressed (Muramatsu, 2011; Sakamoto and Kadomatsu, 2012). Müller glia share common features of quiescence, self renewal and multipotency with cancer stem cells. In unlesioned retina, Muller glia are quiescent and sporadically divide and produce fate-restricted rod precursor (Raymond and Rivlin, 1987; Stenkamp, 2011). Cell death reprograms Müller glia to a stem cell-like state(Goldman, 2014; Gorsuch and Hyde, 2014; Hamon et al., 2016; Karl and Reh, 2010; Lenkowski and Raymond, 2014). *In vitro* experiments demonstrated that inhibition of Midkine successfully suppresses proliferation of cancer stem cells(Erdogan et al., 2017; Mirkin et al., 2005). Therefore, Midkine silencing is proposed as a potential therapy for limiting cell cycle progression in cancer stem cells(Muramatsu and Kadomatsu, 2014). Our data also provide molecular insights into the potential role of Midkine in tumorigenesis, especially in regulating the cell cycle among cancer stem cells.

Constitutive activation of glial cells and formation of a glial scar are detrimental to the function of the central nervous system. An intriguing phenotype in the *mdka*^*mi5001*^ mutants is the cell death-induced gliotic remodeling of Müller glia. A previous report showed that pharmacological suppression of cell cycle progression following photoreceptor death results in hypertrophy and increased GFAP in Müller glia(Thomas et al., 2016). Together, these results suggest that in zebrafish Müller glia the molecular mechanisms that promote cell cycle progression are required to limit the initial gliotic response. Although it is not clear whether entry into the cell cycle initiates reactive gliosis in mammalian retinas, levels of cell cycle proteins appear to be a critical variable in the gliotic response(Dyer and Cepko, 2000; Levine et al., 2000; Ueki et al., 2012; Vázquez-Chona et al., 2011).

Our data significantly expand the understanding of retinal regeneration in zebrafish and more fully define the function of Midkine-a in governing the eukaryotic cell cycle. We provide convincing evidence that Midkine-a regulates proliferation of reprogrammed Müller glial during the regeneration of cone photoreceptors. In the absence of Midkine-a, zebrafish Müller glial respond similarly to Müller glia in mammals, with only a limited ability to regenerate neurons. In developing mammalian retinas, Midkine has been identified as component in the core transcriptional repertoire of mitotic retinal progenitors(Livesey et al., 2004). It remains to be determined if Midkine-dependent cell cycle machinery is present in the Müller glia of adult mammals or if manipulation of Midkine signaling in adult mammals could promote neuronal regeneration.

## Supporting information

Suppl. Table 1; Differential Gene Expression

Suppl. Table 2; Pathway Analysis

Suppl. Table 3; List of Antibodies

Suppl. Table 4; List of Primers

## Acknowledgments

The authors thank Dilip Pawar and Alex LeSage for zebrafish maintenance and technical assistance. This work was supported by grants from the National Institutes of Health (NEI) - R01EY07060 and P30EYO7003 (PFH) and an unrestricted grant from the Research to Prevent Blindness, New York. Fish lines and reagents provided by ZIRC were supported by NIH-NCRR Grant P40 RR01.

## Materials and Methods

### Zebrafish

Fish were maintained at 28C on a 14/10 h light/ dark cycle with standard husbandry procedures. Zebrafish lines were created on the AB-wildtype background. All animal procedures were approved by the Institutional Animal Care and Use Committee at the University of Michigan.

### CRISPR-Cas9 mediated targeted mutation of *midkine-a*

Targeted mutations in the *midkine-a* locus were introduced using CRISPR-Cas9(Hwang et al., 2013). Briefly, ZiFit software (zifit.partners.org) was used to identify guide RNA target sequence for midkine-a (Fig. S1A). Oligos to the target sequencing (GGC AGC TGC GTG GCC AAT AAC GG) were annealed and subcloned into the pT7 gRNA vector(Hwang et al., 2013; Jao et al., 2013; Nasevicius and Ekker, 2000). The subcloned vector was digested with BamHI and sgRNA was transcribed with MEGAshortscript T7 kit (Thermo Fisher Scientific, Waltham, MA). To synthesize *cas9* mRNA, pCS2nCas9n plasmid and mMessage mMachine SP6 in vitro transcription kits (Thermo Fisher Scientific) were used. Purification of sgRNA and mRNA was performed using mirVana miRNA isolation kit (Thermo Fisher Scientific) and RNsasy mini Kit (Qiagen, Venlo, Netherlands). Single cell stage embryos were injected with 1 nL solution, containing 150 pg *cas9* mRNA and 100 pg sgRNA diluted in 1XDanieux buffer with 2.5% phenol red. F0 embryos were raised to adulthood and then outcrossed with AB-wildtype animals. To screen potential mutants in F1 generation, genomic DNA fragment containing the midkine-a target site was amplified with primers (F; TGACTTTGAAGCTTATTGACGCTG; R: GTGCAGGGTTTGGTCACAGA) and was subjected to T7 endonuclease assay. PCR products with potential indel mutation in the midkine-a gene were sequenced and analyzed with National Center for Biotechnology Information (NCBI) Basic Local Alignment Search Tool and ExPaSy translate tool (www.expasy.org). F1 progenies with indel mutation were in-crossed and homozygous F2 mutants were identified.

### Western Blots

Western Blot analyses were performed as previously described(Calinescu et al., 2009b). Briefly, proteins were extracted from the heads of 30-50 wildtype and *mdka*^*mi5001*^ embryos in cold RIPA lysis buffer containing protease and phosphatase inhibitor cocktail (Cell Signaling Tec., Danvers, MA). Proteins were separated in 12%Mini-PROTEIN TGX Precast gel (BioRad, Hercules, CA) and were transferred to polyvinylidene difluoride (PVDF) membranes (GenHunter Corp., Nashville, TN). After blocking in 5% nonfat dry milk in Tris-buffered saline containing 0.3% Tween-20, membranes were incubated with rabbit anti-Midkine-a antisera followed by horseradish peroxidase-conjugated secondary antibody (1:1000)(Calinescu et al., 2009b). Immunolabeled proteins were detected using the enhanced ECL detection system for chemiluminescence assay (Amersham Biosciences, Arlington Heights, IL). Actin was used as a loading control.

### RNAseq

Embryos at 30 hpf were manually dechlorinated. De-yolking was performed by triturating with glass pipette in cold Ringer’s solution containing 1mM EDTA and 0.3mM phenylmethylsulfonylfluoride in isopropanol. Total RNA from 30 embryos was extracted using TRIzol (invitrogen, Carlsbad, CA). Purity of RNA was analyzed with Bioanalyzer (Agilent Technologies, Santa Clara, CA). Samples with an RNA integrity number of acceptable quality (>7) were used for Illumina RNA-seq library preparation. Deep sequencing was performed on an Illumina GAIIx sequencer (Illumina, Inc., San Diego, CA).

### Read quality trimming and quality assessments

Trim Galore! (v0.2.7; Babraham Institute, Cambridge UK) was used to trim adapter sequences and poor quality bases (below Phred of 20) from the reads while removing any reads that were less than 20 nucleotides long, using the default parameters. Trim Galore! makes use of cutadapt (v1.4.2) (-f fastq -e 0.1 -q 20 -O 1 -a AGATCGGAAGAGC file.fq.gz). The quality of the reads was assessed before and after trimming with FastQC (v0.10.1).

### Read mapping and gene-level quantitation

Quality trimmed and filtered reads were aligned to release 83 of the GRCz10 Ensembl genome build with bowtie2 (v2.2.6) and gene-level quantitation was performed with RSEM (v1.2.22). This was done using the rsem-calculate-expression command from RSEM, which calls bowtie2 (--sensitive --dpad 0 --gbar 99999999 --mp 1,1 --np 1 --score-min L,0,-0.1) and streams reads into RSEM for quantitation.

### Differential expression analysis and annotation

The gene-level counts output from RSEM were filtered to remove noise prior to normalization with trimmed means of M (TMM), such that only genes with a FPKM (fragments per kilobase of exon per million reads mapped) value greater than 1 in all replicates of any genotype were retained. Counts per million (CPM) were determined using edgeR (v3.10.2), genes with a CPM < 1 in all samples were removed, and remaining counts were TMM normalized. Limma (v3.24.15) was used to voom transform the filtered count data by empirically deriving and applying quality weights to the samples. These weighted values were used to calculate differential expression using limma. Annotations for each gene were added using biomaRt (v2.24.0) including both the *Danio rerio* Entrez gene identifiers and the corresponding *Mus musculus* Entrez orthologous gene identifiers.

### Gene Ontology and Pathway Analysis Overview of workflow

Gene Ontology (GO) term enrichment analysis was performed using a log2-fold change (log2FC) ranked list from limma (log2FC > 1 and FDR < 0.05) as input into clusterProfiler (v2.2.4). This analysis determines which Molecular Function, Biological Process, or Cellular Component GO terms are positively or negatively enriched in the mutant embryos compared with wild-type, at a false discovery rate ≤ 0.05, while taking into account the magnitude and direction of change. Pathway database, Reactome pathway analyses, were used. A log2FC ranked list of all differential gene expression data from limma (log2FC > 1 and FDR < 0.05) was input into ReactomePA (1.12.3).

All the zebrafish genes in the dataset were manually annotated with their murine orthologs using biomaRt, and a Reactome pathway analyses was performed using zebrafish gene annotations (from zebrafish differential expression data) and zebrafish pathway annotations. The analysis was performed with zebrafish Entrez gene identifiers to determine the Reactome pathways that were positively or negatively enriched at a false discovery rate ≤ 0.05.

### EdU and BrdU labeling

Proliferating cells were labeled with the S-phase markers, 5-ethynyl-2’deoxyuridine (EdU; Thermo Fisher Scientific) or 5-Bromo-2’deoxyuridine (BrdU; Sigma-Aldrich Corp., St. Louis, MO). Embryos were incubated on ice in 1.5 mM EdU dissolved in embryo rearing solution containing 15% diethylsulfoxide for 20 minutes. Embryos were then returned to room temperature for 10 minutes prior to fixation. EdU labelled cells were visualized using Click-iT Assay kit (Thermo Fisher Scientific). To label dividing cells in adults, animals were housed in the fish system water containing 5 mM BrdU and processed as described previously(Gramage et al., 2015).

### Light lesion

To selectively damage photoreceptors, an ultra-high intensity light lesion was used as previously described(Bernardos et al., 2007). In brief, zebrafish were exposed to 120,000 lux light from an EXFO X-Cite 120W metal halide lamp for 30 min and then returned to the aquarium system.

### Histology

Embryos and dissected eyes were fixed in 4% paraformaldehyde with 5% sucrose in PB, pH 7.4 at room temperature for 2 hours or 4C overnight, respectively. After rinsing with 5% sucrose in PBS, tissues were cryoprotected, embedded, and sectioned at 6 µm as previously described (19). Immunocytochemistry was performed as previously described(Bernardos et al., 2007). Primary and secondary antibodies used in this study was listed in Supplementary Table 1.

### Flat-mount retinal immunocytochemistry

Retinas were isolated from the eyes of dark-adapted zebrafish and fixed overnight at 4C in 4% paraformaldehyde in PB with 5% sucrose, pH 7.4. Immunocytochemistry was performed as previously described(Nagashima et al., 2017).

### Microscopy, image analysis, and cell counts

Retinal cross sections and flat-mount retinas were imaged with Leica DM6000 Upright Microscope System and Leica TCS SP5 confocal microscope (Leica Microsystems, Wetzlar, Germany), respectively. Adobe Photoshop CS6 Extended (Adobe Systems, San Jose, CA), Leica Application Suite X (Leica Microsystems), ImageJ (https://imagej.nih.gov/ij/), or icy (http://icy.bioimageanalysis.org/) were used for image analysis, 3D reconstruction, and movie production. ZO1 profiles with perimeter greater than 3.5 µm were identified as cone photoreceptors. For each retina, 6 areas were sampled, 5625 µm^2^/µm total area, and cones were counted from confocal images using NIH ImageJ (https://imagej.nih.gov/ij/). Statistical analysis was performed in JMP pro software (SAS Institute Inc., Cary NC).

### qPCR

Total RNA from whole retinas was extracted using TRIzol (6 retinas from 3 fish per sample) (Invitrogen). RNA was quantified using Nanodrop spectrophotometer (Thermo Fisher Scientific). Reverse transcription and qPCR were performed according to the manufacturer’s instructions using Qiagen QuantiTec Reverse Transcription kit and BioRad IQ SYBR Green Supermix, respectively. Reactions were performed using a CFX384 Touch Real-Time PCR Detection Systems (Bio-Rad). Three biological replicates were prepared for each time point, and three technical replicates were evaluated for each sample. For quantification of fold changes, ΔΔ C(t) method was used, and the housekeeping gene, *gpia*, was used to normalize the data(Livak and Schmittgen, 2001). Primer sequences are listed in the supporting table 2. Statistical comparison were performed using JMP pro.

### ALK Inhibitor treatment

Zebrafish were housed from 24 to 72 hpl in system water containing 10 µM TAE684 (Abcam, ab142082, Cambridge, UK) constructed from a 20 mM stock solution in 0.1% dimethylsulfoxide. Control groups were housed in system water containing the 0.1% dimethylsulfoxide. Solutions were changed daily.

## Supplemental Information

**Fig. S1.**
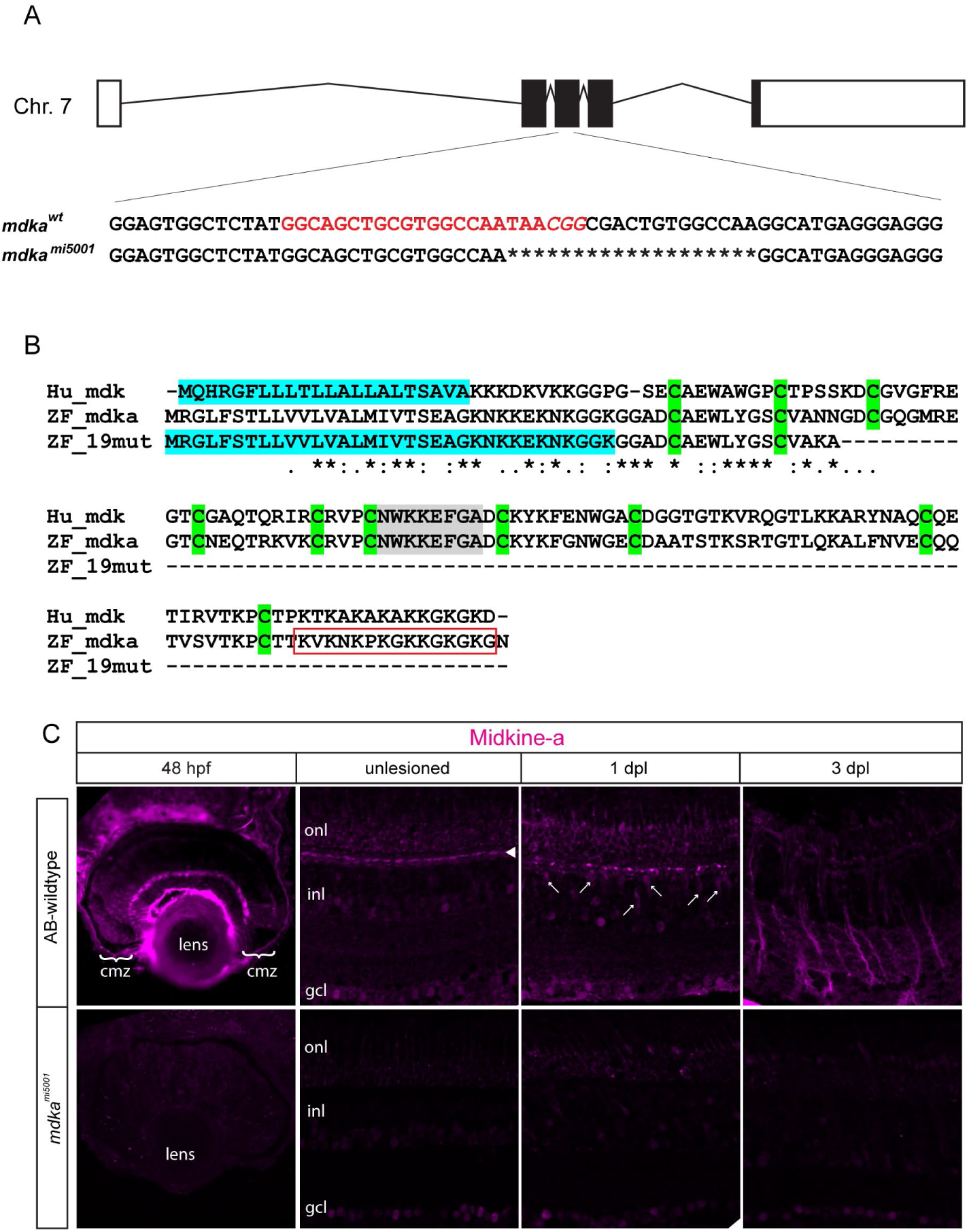
CRISPR-Cas9 mediated mutation in the *midkine-a* locus. Schematic representation of the *midkine-a* locus on Chromosome 7. The gene consists of five exons. The gRNA target sequence located at exon 3 is shown in red. (B) Protein sequence alignment from human MIDKINE and zebrafish Midkine-a from wildtype and mutant animals. The 19 bp deletion introduces a premature stop codon, resulting in a truncated protein. (C) Immunocytochemistry for Midkine-a in embryonic (48 hpf), adult, and regenerating retinas. In unlesioned retina, Midkine-a immunoreactivity is detected in horizontal cells (arrowhead and(Gramage et al., 2015)). Photoreceptor cell death induces Midkine-a in reprogrammed Müller glia (arrows) at 1 dpl. Midkine-a distributes radial processes of Müller glia at 3 dpl. opl: outer plexiform layer; inl: inner nuclear layer; ipl: inner plexiform layer; gcl: ganglion cell layer. Scale bars; 50 μm.

**Fig. S2.**
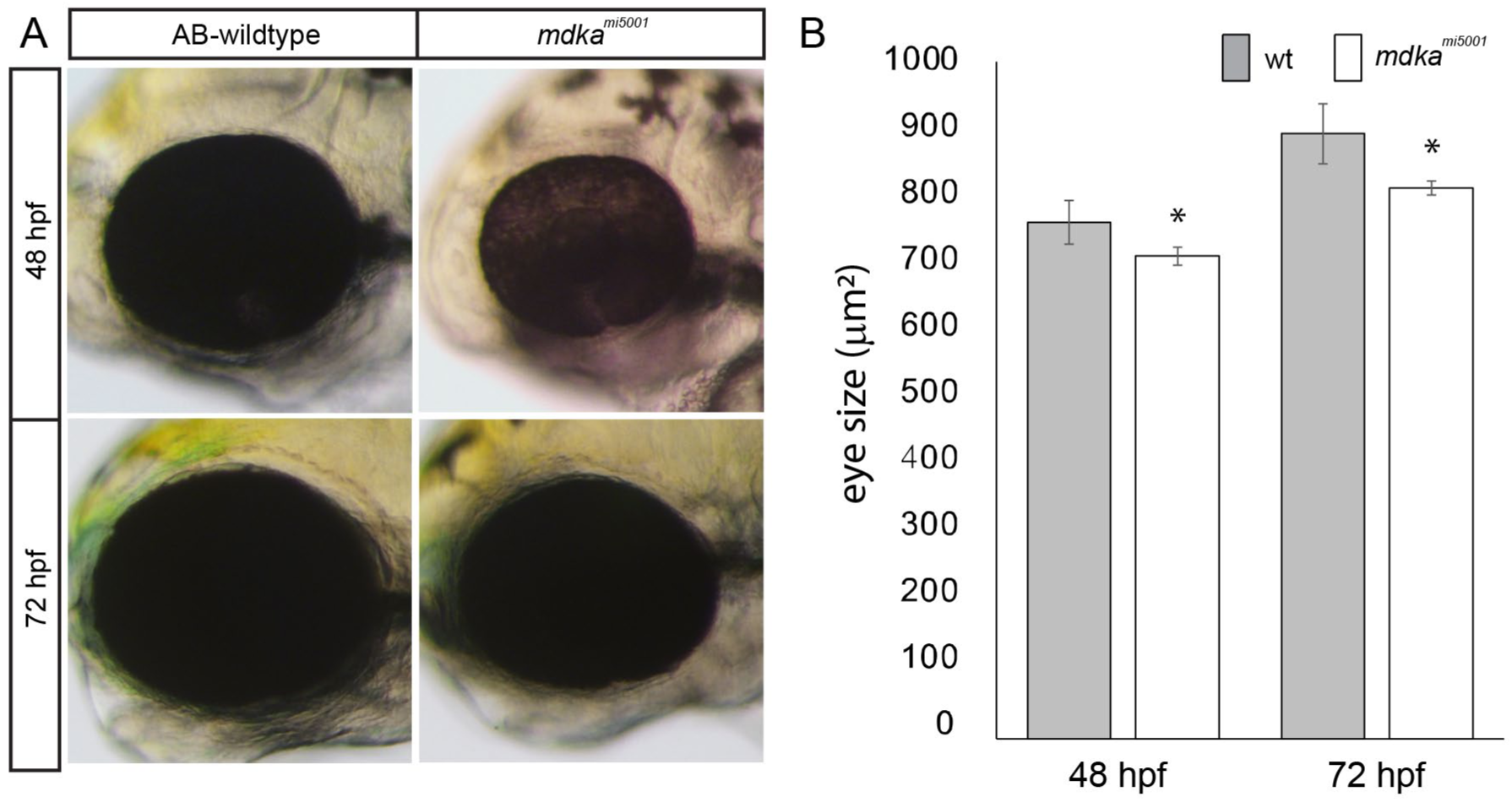
Comparisons of eye size during development. (A-B) Comparison of eye size between wildtype and *mdka*^*mi5001*^ mutant at 48 and 72 hours post fertilization (hpf). The *mdka*^*mi5001*^ has smaller eyes at 48 and 72 hpl. *p<0.01.

**Fig. S3.**
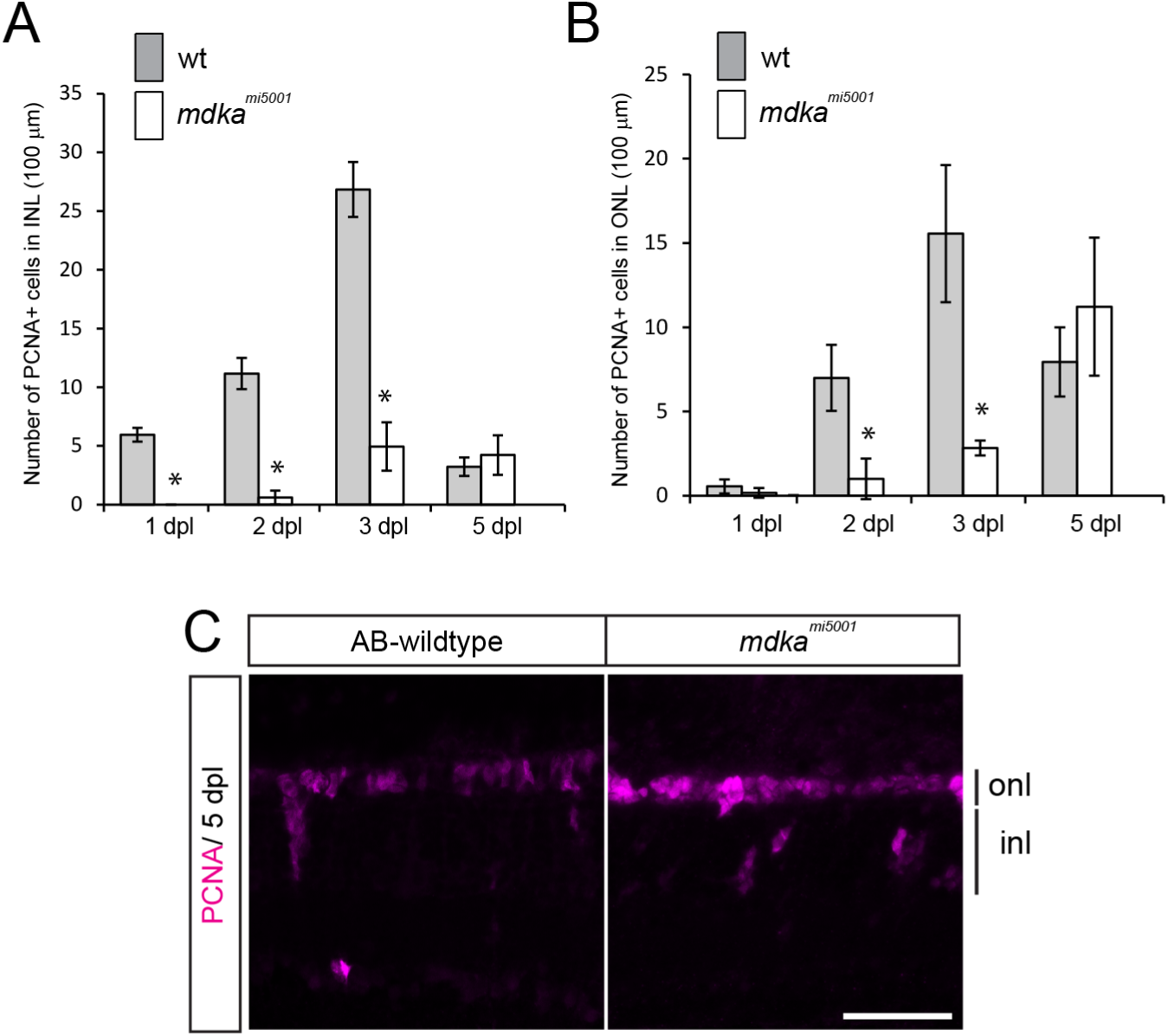
Quantitation of PCNA-positive cells in wildtype and *mdka*^*mi5001*^ retinas. (A-B) The number of PCNA-positive cells in wildtype and *mdka*^*mi5001*^ in the inner (A) and outer (B) nuclear layers. The *mdka*^*mi5001*^ retinas have significantly less PCNA-positive cells at 1, 2, and 3 days post lesion (dpl) compared with wildtype retinas. *p<0.01. (C) Immunocytochemistry for PCNA (magenta) at 5 dpl. PCNA-positive cells mainly localize in the ONL, where rod precursors proliferate, both in wildtype and *mdka*^*mi5001*^ retinas. Scale bar; 30 μm (A). onl: outer nuclear layer; inl: inner nuclear layer.

**Fig. S4.**
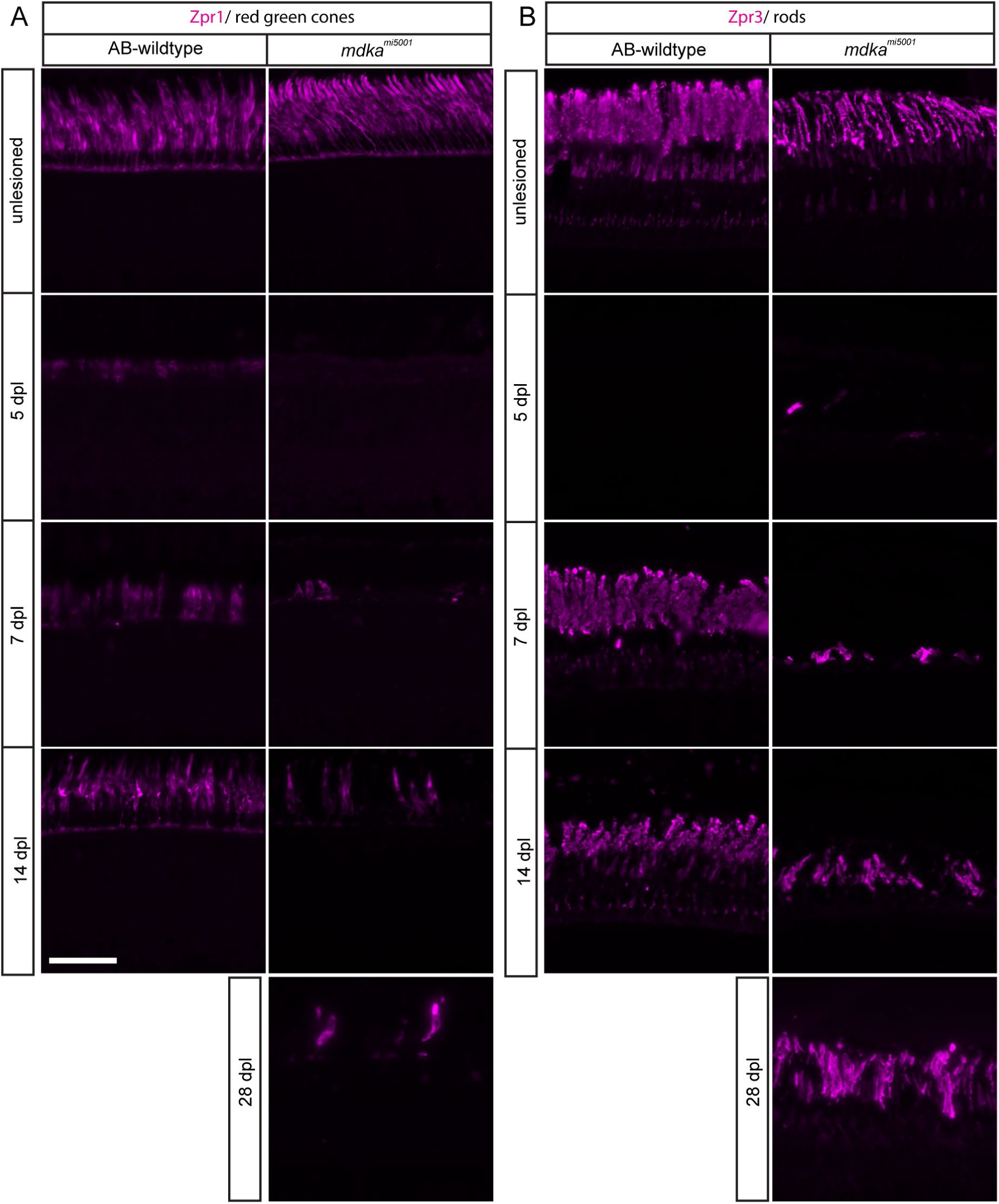
Time course of cone and rod photoreceptor regeneration in wildtype and *mdka*^*mi5001*^ retinas. (A) Immunocytochemistry for red/ green cone photoreceptor marker, Zpr1 (magenta). In wildtype retina, immature cone photoreceptor start to appear at 5 days post lesion (dpl) and regeneration largely completes by 14 dpl. In the *mdka*^*mi5001*^ mutant, regenerating photoreceptors are absent at 5 dpl. At 7 dpl, very few cone photoreceptors appear. The number of cone photoreceptors is less at 14 and 28 dpl compared with wildtype. (B) Immunocytochemistry for rod photoreceptor marker, Zpr3 (magenta) following lesion. In wildytpe retina, regenerating rod photoreceptors appear by 7 dpl. In the *mdka*^*mi5001*^ retinas, rod photoreceptors slowly regenerate. Scale bar; 30 μm.

**Fig. S5.**
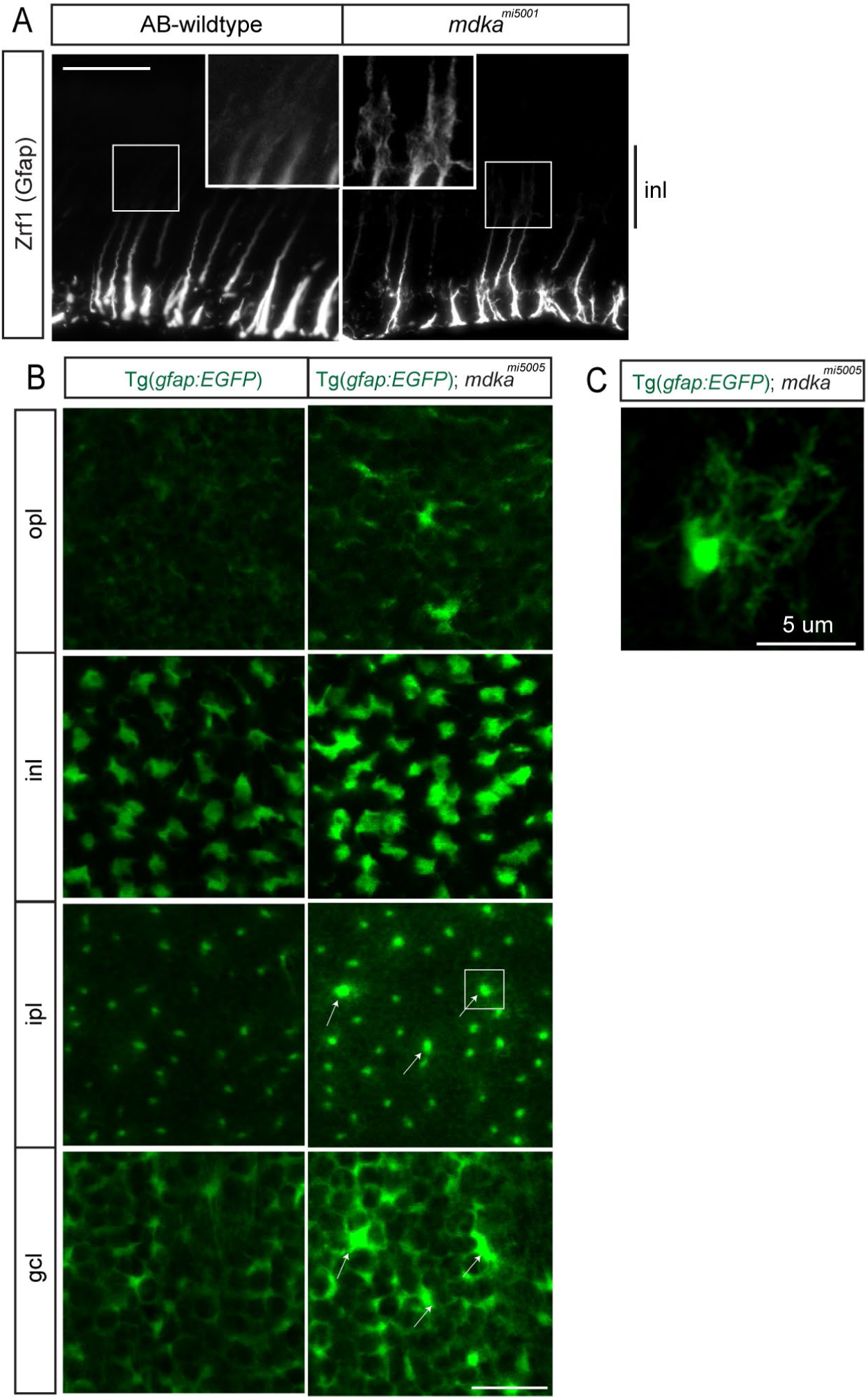
Gliotic remodeling of Müller glia in the *mdka*^*mi5001*^ retinas. (A) Immunocytochemistry for Gfap (white) in wildtype and *mdka*^*mi5001*^ retinas at 28 days post lesion (dpl). In wildtype, the Gfap immunosignal is restricted to the inner third of radial processes. No obvious signal is detected at the inner nuclear layer. The *mdka*^*mi5001*^ upregulates Gfap and signals are seen at the cell body of Müller glia in the inner nuclear layer. (B) Single optical planes from Z-stack series of the Tg(gfap: EGFP) reporter flat-mount retinal preparation in the *mdka*^*mi5001*^ background. In the ganglion cell and inner plexiform layers, some Müller glia show signs of hypertrophy, including increased levels of the EGFP transgene signal (arrows). (C) Higher magnification view of hypertrophic Müller glia (boxed region in B). The hypertrophic Müller glia have expanded lateral processes. Scale bars; 30 μm (A) : 20 μm (B): 5 μm (C). opl: outer plexiform layer; inl: inner nuclear layer; ipl: inner plexiform layer; gcl: ganglion cell layer.

**Fig. S6.**
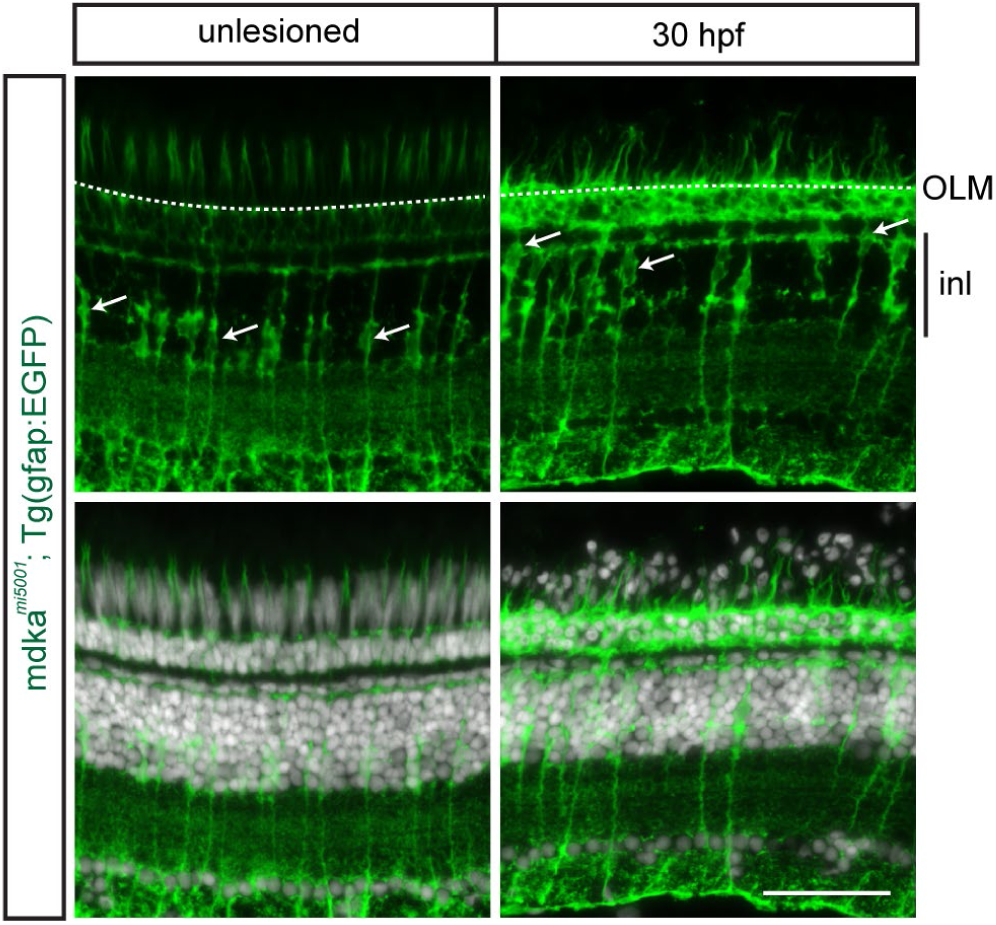
Müller glia in the *mdka*^*mi5001*^ undergo normal interkinetic nuclear migration following photoreceptor death. The *mdka*^*mi5001*^ retinas carrying the Müller glial reporter, Tg(gfap: EGFP). Photoreceptor injury induces interkinetic nuclear migration of Müller glial nuclei in the *mdka*^*mi5001*^ mutant. Cell bodies of Müller glia are indicated by arrows. Scale bar; 30 μm. OLM: outer limiting membrane; inl: inner nuclear layer.

**Fig. S7.**
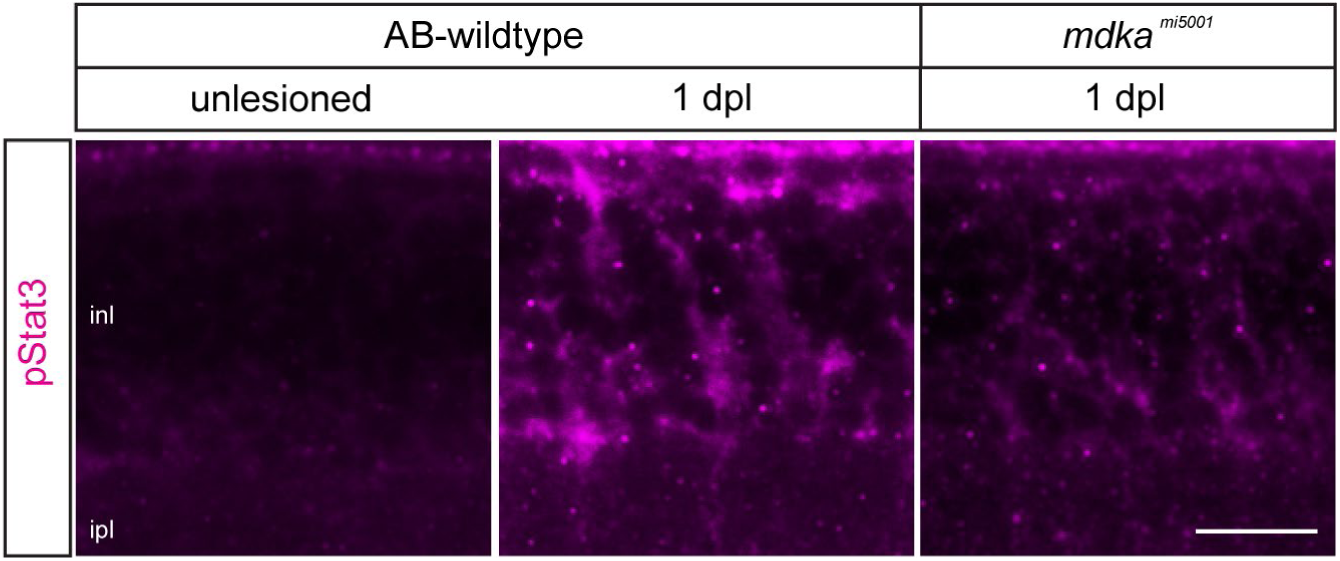
Stat3 phosphorylation is diminished in the *mdka*^*mi5001*^ mutant. Immunocytochemistry for phosphorylated Stat3 (pStat3) in the wildtype and *mdka*^*mi5001*^ mutant. In unlesioned retina, immuno-signal for pStat3 is not detected in the inner nuclear layer. Following photoreceptor lesion, Müller glia in the inner nuclear layer upregulate pStat3 in wildtype, whereas *mdka*^*mi5001*^ mutants have reduced phosphorylation of Stat3. Scale bar; 30 μm. inl: inner nuclear layer; ipl: inner plexiform layer.

**Movie S1. Confocal z-stack series of flat-mount retina in *mdka***^***mi5001***^ **; Tg(*gfap: EGFP*) mutant.** Müller glia in the *mdka*^*mi5001*^ remain hypertrophic and increased level of the transgene, gfap: EGFP at 28 dpl.

**Movie S2. 3D reconstruction of the Müller glia in the *mdka***^***mi5001***^ **mutant undergo gliotic remodeling.** Müller glial somata migrate into the outer plexiform layer at 28 days post lesion.

**Movie S3. 3D reconstruction of the Müller glia in the *mdka***^***mi5001***^ **mutant undergo gliotic remodeling.** The abnormal gliotic remodeling Müller glia in the *mdka*^*mi5001*^ at 28 days post lesion.

**Additional data Table S1 (separate file)**

Differential gene expression analysis of whole embryos at 30 hours post fertilization.

**Additional data Table S2 (separate file)**

Pathway analysis of whole embryos at 30 hours post fertilization.

