## Supplementary material for "Midkine-a is required for cell cycle progression of Müller glia during neuronal regeneration": Suppl. Table 3; List of Antibodies

Table S3 Antibody list

| antibody description | Source | dilution |
|---|---|---|
| mouse anti-PCNA | Sigma Aldrich: P8825 | 1:1000 |
| mouse anti-Zpr1 | Zebrafish International Resource Center | 1:200 |
| mouse anti-ZO1 | ThermoFisher Scientific: ZO1-1A12 | 1:200 |
| guinea pig anti-Rx1 | Open Biosystems (Nagashima et al., 2013) | 1:200 |
| mouse anti-Sox2 | Genetex: T+GTX124477 | 1:200 |
| mouse anti-BrdU | Abcam: ab6326 | 1:100 |
| rabbit anti-pAlk | Abcam: ab111865 | 1:200 |
| mouse anti-Zpr3 | Zebrafish International Resource Center | 1:200 |
| mouse anti-Zrf1 | Zebrafish International Resource Center | 1:200 |
| mouse anti-pStat3 | MBL: D128-3 | 1:200 |
| chick anti-GFP | Abcam: | 1:400 |
| anti-mouse IgG Alexa Fluor 555 | ThermoFisher Scientific: A21422 | 1:200 |
| anti-mouse IgG Alexa Fluor 633 | ThermoFisher Scientific: A21050 | 1:200 |
| anti-rabbit IgG Alexa Fluor 633 | ThermoFisher Scientific: A21071 | 1:200 |
| anti-rat IgG Alexa Fluor 555 | ThermoFisher Scientific: A21434 | 1:200 |
| anti-chick IgG Alexa Fluor 488 | ThermoFisher Scientific: A11039 | 1:200 |
